# Sex differences in DNA demethylation machinery precede sex differences in the oxytocinergic system in the postnatal mouse brain

**DOI:** 10.64898/2026.08.10.744005

**Authors:** R Bigarani, B Ghione, MJ Cambiasso, CD Cisternas

**Author notes:** Corresponding author: Dr. Carla Daniela Cisternas. Email of the authors: Rocío Bigarani, Brunella Ghione, María Julia Cambiasso.

## Abstract

In mammals, sex differences in the brain arise from genetic and hormonal factors, including organizational effects of perinatal testosterone. Epigenetic mechanisms including DNA methylation and demethylation have emerged as critical mediators of brain masculinization; specifically, their regulatory enzymes are upregulated in neonatal mice during the critical period of sexual differentiation, with their inhibition abolishing sex-specific cellular phenotypes. Here, we assessed sex differences in gene expression of the DNA demethylation machinery (*Tet1*, *Tet2*, *Tet3*, *Gadd45a*, *Gadd45b* and *Tdg*) during and after the critical period, and examined how these differences relate to the oxytocinergic system. mRNA expression was measured in the prefrontal cortex (PFC), preoptic area (POA) and paraventricular nucleus of the hypothalamus (PVN) at postnatal day (P) 7 and P18. At P7, males showed higher expression of all six genes than females in PFC, with no differences in POA or PVN; by P18, no regional differences remained. Oxytocin (OXT) immunoreactivity was surveyed across periventricular nucleus (Pe), anteroventral periventricular nucleus (AVPe), POA, PVN and supraoptic nucleus (SON). OXT was undetectable in the POA, AVPe and Pe at P7, and no sex differences were found in PVN or SON at either age, or in AVPe at P18. At P18, females showed higher OXT-immunoreactivity in the Pe and POA, than males. For *Oxtr*, qPCR revealed higher mRNA expression in the PFC of males at P7, with no other regional differences and none remaining at P18. Together, these findings suggest that sex differences in oxytocinergic regions arise from sex-specific epigenetic regulation during the critical period, and that perinatal testosterone may program DNA methylation dynamics underlying sex-specific gene expression in the developing brain. Our results support a model in which testosterone-dependent epigenetic mechanisms contribute to the sexual differentiation of neuroendocrine circuits, linking hormonal signals to long-term brain organization.

## 1. INTRODUCTION

A large proportion of the sex differences described in the brain arise from a transient perinatal exposure to gonadal testosterone in males, during an early developmental window known as the critical period of brain sexual differentiation, which in rodents spans from late gestation to approximately postnatal day (PN) 10 (Wilson & Davies, 2007). The classical hypothesis of brain sexual differentiation holds that gonadal steroids organize male-typical neural circuits during this early period, whereas in females the absence of an equivalent hormonal environment results in a default developmental pattern (Wilson & Davies, 2007). The effects of testosterone, or its metabolite 17β-estradiol (E2), are in many cases permanent despite this transient exposure, leading to the concept of a cellular memory of early hormone action. A growing body of evidence indicates that epigenetic mechanisms, particularly DNA methylation, contribute to establishing this cellular memory during development (Mosley et al., 2017; Murray et al., 2009; Nugent et al., 2015).

DNA methylation (5-methylcytosine, 5mC), catalyzed by DNA methyltransferases (DNMTs), was long considered a stable, repressive epigenetic mark, with passive dilution during replication as its only means of removal. This view was challenged by evidence that 5mC levels in the mammalian brain are dynamic, particularly during development (Lister et al., 2013). The discovery of the ten-eleven translocation (TET) enzymes, which initiate active DNA demethylation by converting 5mC to 5-hydroxymethylcytosine (5hmC), opened an additional route of epigenetic regulation (Kriaucionis & Heintz, 2009) whose relevance to the sexual differentiation of the brain has only begun to be explored. Our group previously showed that the expression and activity of the enzymes that place (DNMTs) and remove (TETs) methylation marks peak during the first postnatal week in the mouse hypothalamus and hippocampus, coinciding with the critical period of sexual differentiation, and that there are region-specific sex differences in their expression: males show greater expression of Tet2 and Tet3 in hypothalamic regions during this period, whereas females show greater Tet expression in the hippocampus (Cisternas, Cortes, Bruggeman, et al., 2020). Moreover, neonatal inhibition of DNA methylation disrupts masculinization of neurochemical phenotypes assessed later in life (Cisternas, Cortes, Golynker, et al., 2020), and neonatal knockdown of Tet2/Tet3 is sufficient to feminize estrogen receptor alpha (ERα) expression in the male arcuate nucleus (Cortes et al., 2022). Together, these findings suggest that both methylation and active demethylation contribute, in a region-and gene-specific manner, to the establishment of sex differences in neurochemical phenotype (Cortes & Forger, 2023).

Active DNA demethylation is not limited to the oxidative conversion mediated by TET enzymes: 5hmC and its further oxidized derivatives (5-formylcytosine, 5-carboxylcytosine) can be recognized by a base-excision repair (BER) pathway, in which Gadd45 (growth arrest and DNA damage-inducible 45) proteins recruit the repair machinery to specific genomic sites, with thymine DNA glycosylase (TDG) ultimately removing the modified cytosine (Li et al., 2015; Niehrs & Schäfer, 2012). Evidence indicates that this portion of the demethylation pathway is also sexually differentiated during development: in the neonatal rat dentate gyrus, the greater neurogenesis observed in males has been linked to lower *Gadd45a* expression and, consequently, more limited demethylation in that sex and region (Stockman et al., 2022). However, whether a similar pattern of sex-specific regulation of *Gadd45* and *Tdg* occurs in other brain regions, such as the prefrontal cortex, and whether it coincides temporally with TET enzyme expression during the critical period, remains unknown.

Among the neurochemical systems potentially regulated by these epigenetic mechanisms is the oxytocinergic system. Oxytocin (OXT)-producing neurons are located mainly in the paraventricular (PVN) and supraoptic (SON) hypothalamic nuclei, although cell populations have also been described in the preoptic area (POA), the periventricular nucleus (Pe), the anteroventral periventricular nucleus (AVPe), and the bed nucleus of the stria terminalis (BNST) (Madrigal & Jurado, 2021). The distribution and number of OXT+ neurons change markedly across postnatal development, with progressive consolidation of the different nuclei occurring between birth and the third postnatal week (Madrigal & Jurado, 2021; Soumier et al., 2022). The oxytocin receptor (OXTR, *Oxtr*) likewise shows a dynamic, transient pattern of expression during postnatal development, with sex differences described in some subcortical and hypothalamic regions after the second week of life (Newmaster et al., 2020) and in the preoptic area (Sharma et al., 2019) in adult mice, suggesting that the sex-specific organization of the oxytocinergic system depends, at least in part, on perinatal hormone exposure (Dumais et al., 2013).

Despite this evidence, no study to date has jointly characterized, in the same model and during the critical period of brain sexual differentiation, the expression of the active DNA demethylation machinery (*Tet1-3*, *Gadd45a*, *Gadd45b* and *Tdg*) and that of the oxytocinergic system (OXT and *Oxtr*). Such a characterization is a necessary step toward testing the hypothesis that active DNA demethylation contributes to the sexual organization of neurochemical systems regulated by perinatal hormones. Here, we examined, in male and female C57BL6 mice at two postnatal ages: PN7, within the critical period of sexual differentiation, and PN18, after this period has ended, the mRNA expression of *Tet1*, *Tet2*, *Tet3*, *Gadd45a*, *Gadd45b*, *Tdg* and *Oxtr* by qPCR in the prefrontal cortex, preoptic area and paraventricular nucleus, and the protein expression of OXT by immunohistochemistry in the periventricular nucleus, anteroventral periventricular nucleus, preoptic area, paraventricular nucleus and supraoptic nucleus. Our aim was to determine whether sex differences exist in these components and whether their temporal emergence coincides with the critical period, which would support perinatal testosterone as a common modulator of both systems.

## 2. Materials and Methods

### 2.1 Animals

Wild-type C57BL/6 mice were bred and housed in the animal facility of the Instituto de Investigación Médica Mercedes y Martín Ferreyra (INIMEC-CONICET-UNC) under controlled conditions (23 °C, 12:12 h light-dark cycle) with food and water available *ad libitum*. Sex was determined by anogenital distance at the time of tissue collection and subsequently confirmed by gonadal dissection and examination at sacrifice. All procedures were approved by the Institutional Committee for the Care and Use of Experimental Animals (CICUAL) of Instituto Ferreyra (Institutional Resolution 1/2020) and conducted in accordance with national and international guidelines for the care and use of laboratory animals.

### 2.2 Tissue collection

Brains from male and female mice at PN7 and PN18 were rapidly dissected, flash-frozen by immersion in isopentane cooled on dry ice, and stored at −80 °C until use. Coronal sections were obtained in a cryostat maintained at −25 °C, and bilateral tissue punches were collected from the prefrontal cortex (PFC), preoptic area (POA), and paraventricular nucleus (PVN). Punch diameter was 1 mm for all regions and ages, except for the POA at PN7, for which a 0.8 mm punch was used. For tissue punch collection, anatomical landmarks were identified according to the *Atlas of the Developing Mouse Brain at E17.5, P0 and P6* (Paxinos et al., 2007), used as the closest available reference for PN7 tissue, with atlas plates 106–115 for the PFC, 121–124 for the POA, and 125–131 for the PVN. For PN18, anatomical landmarks were identified according to Paxinos and Franklin’s *The Mouse Brain in Stereotaxic Coordinates* (Paxinos & Franklin, 2019), using atlas plates 6–23 for the PFC, 26–33 for the POA, and 33–41 for the PVN. To preserve RNA integrity, the entire procedure was performed within the cryostat chamber.

### 2.3 Reverse transcription and quantitative real time PCR (RT-qPCR)

Tissue samples were homogenized in TRIzol reagent (Invitrogen, USA) by repeated passage through a 30-gauge needle at 4 °C. Total RNA was isolated, and its concentration and purity were assessed by spectrophotometry using a NanoDrop 2000 (Thermo Fisher Scientific, USA). Subsequently, 1 μg of total RNA from each sample was reverse transcribed into cDNA in a final volume of 20 μL using M-MLV reverse transcriptase (Promega, USA) and random primers (Invitrogen, USA), according to the manufacturer’s instructions. Reverse transcription was performed at 37 °C for 60 min, followed by enzyme inactivation at 95 °C for 5 min. Quantitative real-time PCR (qPCR) assays were performed using QuantStudio 3 Real-Time PCR Systems (Applied Biosystems, USA) and FastStart Universal SYBR Green Master Mix (Roche), following a thermal cycling protocol consisting of an initial hold stage of 95 °C for 10 min, followed by 40 cycles of 95 °C for 15 s and 60 °C for 1 min. A melting curve stage (95 °C for 15 s, 60 °C for 1 min, and 95 °C for 1 s) was included at the end of each run to confirm amplification specificity. Primer pairs (Table 1) were designed using Primer-BLAST (National Institutes of Health, USA), selecting sequences spanning exon–exon junctions to ensure mRNA-specific amplification. Primer performance was assessed by standard curve analysis; amplification efficiencies (E = 10–1/slope) ranged between 1.9 and 2.1, corresponding to 90–110%. Specificity was confirmed by melting curve analysis, which showed a single PCR product for each primer pair. Relative gene expression was calculated using the Pfaffl method (Pfaffl, 2001), normalizing to 18S and referencing values to the mean expression of male samples at each postnatal age.

**Table 1.**
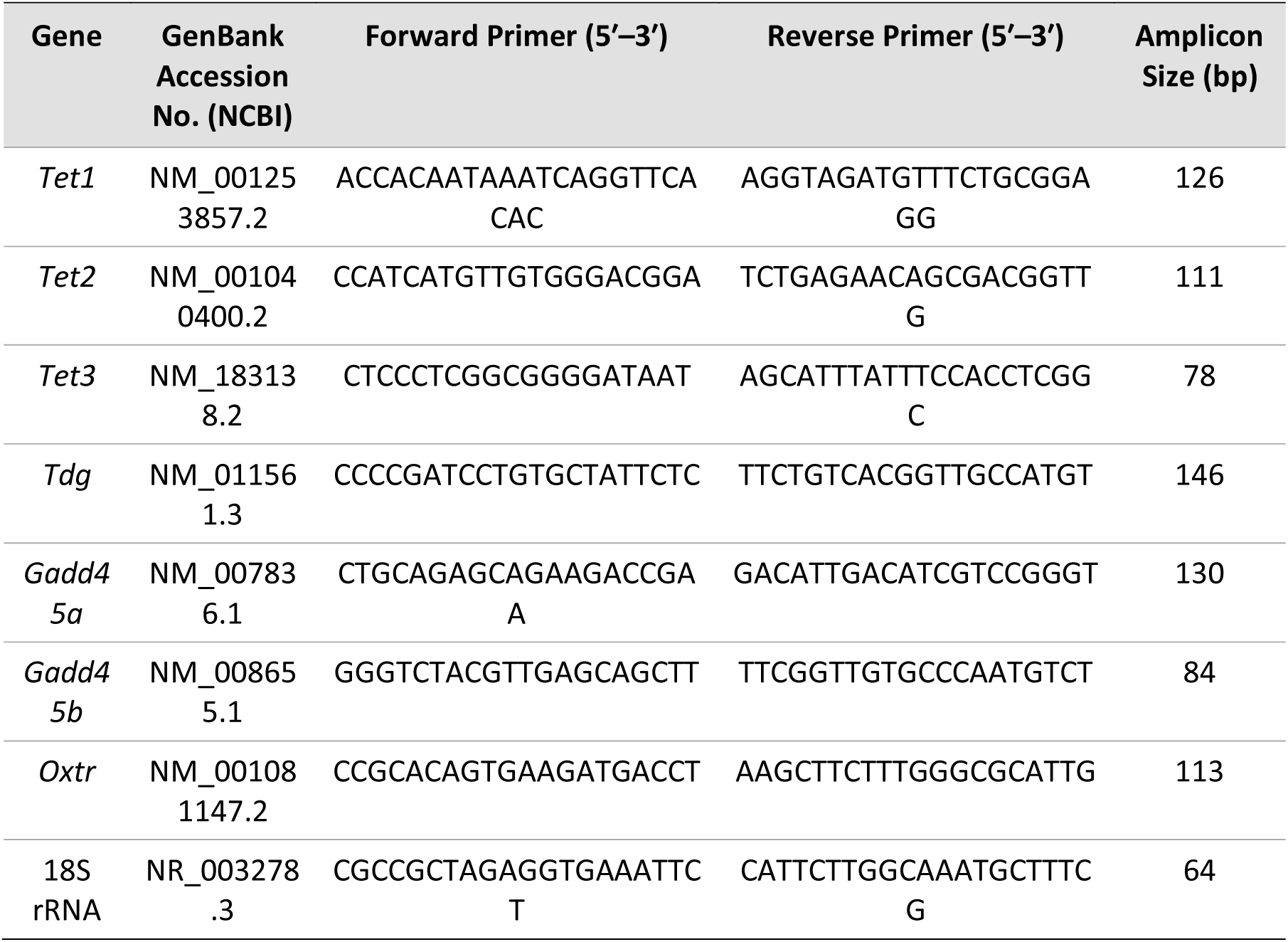
Primer sequences used for qPCR assays.

### 2.4 Immunohistochemistry for oxytocin

Mice were deeply anesthetized by intraperitoneal injection of 5% chloral hydrate (1 g/kg for PN7 mice and 1.4 g/kg for PN18 mice) and transcardially perfused with a washing solution (0.14 M sodium chloride, 0.02 M sucrose, 0.02 M glucose, and 0.2 mL heparin per 100 mL of solution) followed by 4% paraformaldehyde (PFA) in borate buffer (pH 7.4). Brains were then removed, post-fixed in 4% PFA at 4 °C for 24 hours, and cryoprotected in 30% sucrose until sectioning. Coronal brain sections (40 μm) were obtained using a freezing microtome. Each brain was collected into two series, one of which was processed for oxytocin immunohistochemistry. One series of sections was washed in TBS (Tris Buffer Solution: 0.02 M Tris(hydroxymethyl)aminomethane, 0.15 M sodium chloride, pH 7.6), incubated for 30 min in a blocking solution (1X TBS, 10% normal horse serum, 1% hydrogen peroxide, and 0.4% Triton X-100), and then incubated overnight at 4 °C with a rabbit anti-oxytocin primary antibody (1:10,000; Peninsula Laboratories, T4084). The following day, sections were incubated for 1 h at room temperature (RT) with a biotinylated donkey anti-rabbit secondary antibody (1:500; Jackson ImmunoResearch Laboratories), followed by 1 hour incubation at RT with avidin–biotin complex (Vectastain® ABC kit, Vector Laboratories). Immunoreactivity was visualized using 0.05% 3,3′-diaminobenzidine (DAB) in 1X TBS containing 0.01% hydrogen peroxide (H2O2).

### 2.5 Image acquisition and quantification

Oxytocin immunoreactivity was analyzed in brain sections from male and female mice at PN7 and PN18. Oxytocin-immunoreactive nuclei were identified according to the reference atlases for PN7 (Paxinos et al., 2007) and PN18 (Paxinos & Franklin, 2019) across five regions of interest: the periventricular nucleus (Pe), anteroventral periventricular nucleus of the hypothalamus (AVPe), POA, PVN, and supraoptic nucleus (SON). Sections were imaged using a Zeiss microscope equipped with a Leica DC 200 digital camera. A 10x objective was used to image sections containing the Pe, AVPe, POA, and PVN, while a 20x objective was used for the SON. Image acquisition and analysis was performed by an experimenter blind to the sex of the animals, using ImageJ software (version 1.53). The total number of OXT-immunoreactive (OXT+) somata was counted in each section, and the average number of OXT+ cells per animal was calculated by dividing the total count by the number of sections analyzed per region. OXT+ fiber density was assessed as the percentage of thresholded area within a region of interest defined for each section; images were converted to grayscale, and the same threshold was applied across all individuals. Area measurements from individual sections were averaged to generate a single value per animal. The quantification was done in three to four serial coronal sections for each region.

### 2.6 Statistical analysis

Statistical analyses were performed using GraphPad Prism 8.0.1 (GraphPad Software, USA). Group data are presented as mean ± SEM. Normality of the data distribution was assessed using the Shapiro-Wilk test, and homogeneity of variance was evaluated prior to analysis. Group comparisons were performed using Student’s t-test, with Welch’s correction applied when the assumption of equal variances was not met. Differences were considered statistically significant at P < 0.05.

## 3. Results

### 3.1 Expression of active DNA demethylation and DNA repair genes during the critical period of brain sexual differentiation

We first used quantitative RT-PCR to examine the relative expression of genes involved in the active DNA demethylation (*Tet1*, *Tet2*, and *Tet3*) and DNA repair machinery (*Gadd45a*, *Gadd45b*, and *Tdg*) during the critical period of sexual brain differentiation. Gene expression was analyzed in micropunches of PFC, POA, and PVN collected from PN7 male and female mice.

In the PFC, significant sex differences were observed in the relative mRNA expression of all examined demethylation and DNA repair genes at PN7 (Fig. 1, Welch’s t-test: t_Tet1_ = 3.745, P = 0.010, df = 5.890; t_Tet2_ = 3.443, P = 0.015, df = 5.717; t_Tet3_ = 3.197, P = 0.046, df = 3.183; t_Gadd45a_ = 3.474, P = 0.012, df = 6.328; t_Gadd45b_ = 4.300, P = 0.034, df = 2.488; t_Tdg_ = 2.930, P = 0.043, df = 3.966). In all cases, male mice exhibited higher mRNA expression levels than females.

**Figure 1.**
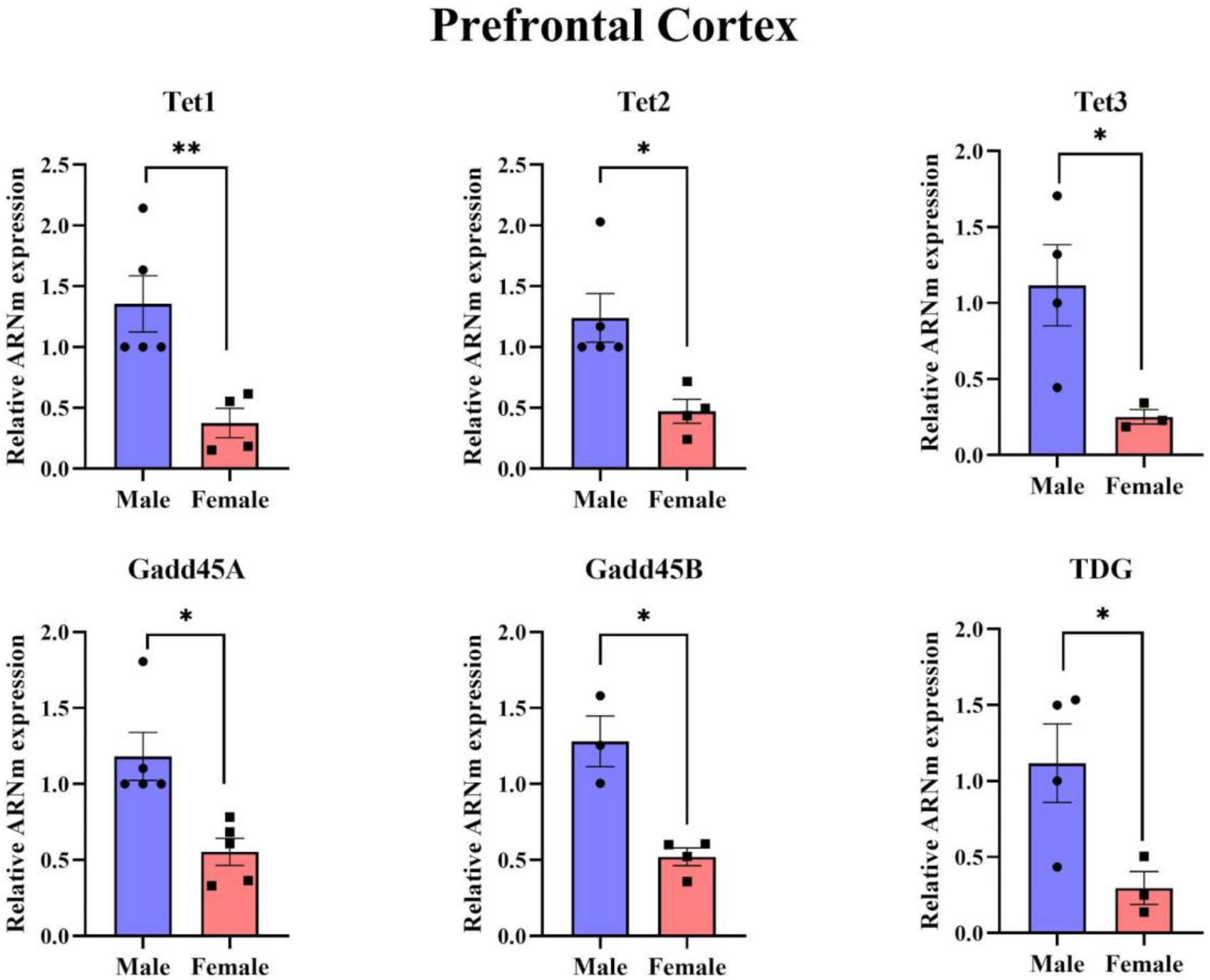
Sex differences in the expression of active DNA demethylation and DNA repair genes in the prefrontal cortex at PN7. Relative mRNA expression of the active DNA demethylation enzymes *Tet1*, *Tet2*, and *Tet3*, and the DNA repair-associated genes *Gadd45a*, *Gadd45b*, and *Tdg* in PFC of male and female mice at PN7. Male mice exhibited significantly higher expression levels of all analyzed genes compared with females. Data are presented as mean ± SEM. Statistical significance was determined using Welch’s *t*-test. *P* < 0.05 was considered statistically significant.

In the POA and PVN, no significant sex differences were detected in the relative mRNA expression of the active DNA demethylation enzymes at PN7 (Suppl. Fig. 1, Welch’s t-test: POA, t_Tet1_ = 0.073, P = 0.944, df = 5.543; t_Tet2_ = 2.245, P = 0.052, df = 8.935; t_Tet3_ = 0.075, P = 0.943, df = 5.775) (Suppl. Fig. 2, Welch’s t-test: PVN, t_Tet1_ = 0.197, P = 0.850, df = 5.914; t_Tet2_ = 0.277, P = 0.798, df = 3.370; t_Tet3_ = 0.366, P = 0.727, df = 6.000). Although no statistically significant differences were observed in these regions, *Tet2* expression in the POA showed a trend toward higher mRNA levels in males compared with females. Likewise, no significant sex differences were found in the expression of the DNA repair-associated genes in either brain region (Suppl. Fig. 1 - 2, Welch’s t-test: POA, t_Gadd45a_ = 0.420, P = 0.699, df = 3.526; t_Gadd45b_ = 0.223, P = 0.830, df = 6.345; t_Tdg_= 0.201, P = 0.851, df = 4.107; PVN, t_Gadd45b_ = 0.454, P = 0.666, df = 5.947; t_Tdg_ = 0.302, P = 0.775, df = 4.971). Notably, *Gadd45a* mRNA expression was not detected in the PVN at PN7.

### 3.2 Expression of active DNA demethylation and DNA repair genes after the critical period of brain sexual differentiation

To determine whether the sex differences observed during the critical period persist after this developmental window, we next examined the relative mRNA expression of the active DNA demethylation enzymes, as well as the DNA repair-associated genes, in the PFC, POA, and PVN at PN18 using RT-qPCR.

No significant sex differences were detected in the relative mRNA expression of *Tet1*, *Tet2*, or *Tet3* in any of the brain regions analyzed (Fig. 2, Welch’s t-test: PFC, t_Tet1_ = 2.074, P = 0.115, df = 3.570; t_Tet2_ = 0.443, P = 0.676, df = 4.950; t_Tet3_ =0.684, P = 0.531, df = 4.023) (Suppl. Fig. 3, Welch’s t-test: POA, t_Tet1_ = 0.152, P = 0.887, df = 3.696; t_Tet2_ = 1.296, P = 0.278, df = 3.282; t_Tet3_ = 0.336, P = 0.755, df = 3.759) (Suppl. Fig. 4, Welch’s t-test: PVN, t_Tet1_ = 1.921, P = 0.105, df = 5.831; t_Tet2_ = 0.498, P = 0.650, df = 3.280; t_Tet3_ = 2.124, P = 0.103, df = 3.864).

**Figure 2.**
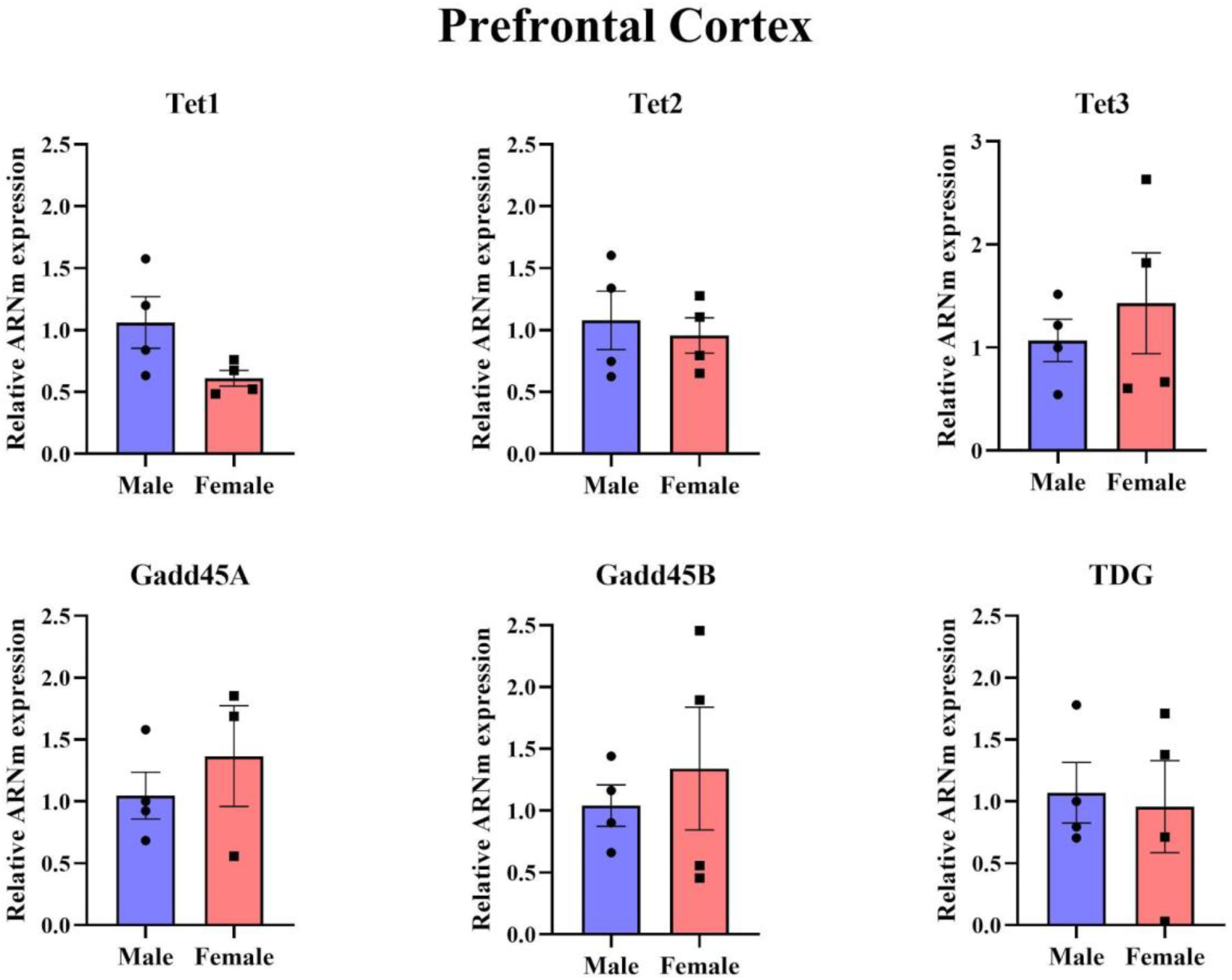
Expression of active DNA demethylation and DNA repair genes in the prefrontal cortex at PN18. Relative mRNA expression of the active DNA demethylation enzymes *Tet1*, *Tet2*, and *Tet3*, and the DNA repair-associated genes *Gadd45a*, *Gadd45b*, and *Tdg* in the PFC of male and female mice at PN18. No significant sex differences were detected for any of the genes analyzed. Data are presented as mean ± SEM. Statistical significance was determined using Welch’s *t*-test. *P* < 0.05 was considered statistically significant.

Furthermore, no significant sex differences were detected in the relative mRNA expression of the DNA repair-associated genes in any of the brain regions analyzed at PN18 (Fig. 2, Welch’s *t*-test: PFC, *t*_Gadd45a_ = 0.709, *P* = 0.532, df = 2.873; *t*_Gadd45b_ = 0.570, *P* = 0.602, df = 3.680; *t*_Tdg_ = 0.252, *P* = 0.811, df = 5.184) (Suppl. Fig. 3, Welch’s t-test: POA, *t*_Gadd45a_ = 0.695, *P* = 0.556, df = 2.117; *t*_Gadd45b_ = 1.146, *P* = 0.346, df = 2.605; *t*_Tdg_ = 1.543, *P* = 0.217, df = 3.174) (Suppl. Fig. 4, Welch’s t-test: PVN, *t*_Gadd45b_ = 0.144, *P* = 0.889, df = 8.959; *t*_Tdg_ = 1.233, *P* = 0.292, df =3.577). As observed during the critical period, *Gadd45a* mRNA expression was not detected in the PVN.

### 3.3 Expression of oxytocin and oxytocin receptor during postnatal brain development

To investigate whether transient early exposure to gonadal hormones contributes to sex differences in the expression of oxytocin (OXT) and its receptor (*Oxtr*) during postnatal brain development, we examined OXT protein levels by immunohistochemistry and quantified *Oxtr* mRNA expression by RT-qPCR in male and female C57BL/6 mice at PN7, corresponding to the critical period of brain sexual differentiation, and at PN18, after the end of this critical period.

As an initial step, a qualitative survey of the entire brain was performed to determine the distribution of OXT immunoreactivity. OXT-immunoreactive cell bodies and fibers were detected in the bed nucleus of the stria terminalis (BNST), the periventricular nucleus (Pe), anteroventral periventricular nucleus (AVPe), POA, PVN, and the supraoptic nucleus (SON). No immunoreactivity was observed in sections processed in the absence of the primary antibody. To facilitate comparison with the results obtained for the active DNA demethylation enzymes and DNA repair-associated genes, quantitative analyses of OXT expression were performed in the same brain regions previously examined: POA, PVN and SON.

#### 3.3.1 Oxytocin immunoreactivity at PN7

At PN7, OXT immunoreactivity was detected in PVN and SON, but it was not detected in the POA, AVPe, or Pe of either sex at this developmental age (Figure 3). Quantitative analysis of the PVN and SON revealed no significant sex differences in either the total number of OXT-immunoreactive neurons or the percentage of OXT-immunoreactive area (Fig. 4A, B). Specifically, in the PVN, the number of OXT+ neurons (Fig. 4A, Welch’s *t*-test: *t* = 0.554, *P* = 0.607, df = 4.398) and OXT-immunoreactive area (Fig. 4A, Welch’s *t*-test: *t* = 1.795, *P* = 0.168, df = 3.090) were comparable between males and females. Similarly, no significant sex differences were observed in the SON for either OXT-positive neuron number (Fig. 4B, Welch’s *t*-test: *t* = 0.761, *P* = 0.481, df = 4.910) or OXT-immunoreactive area (Fig. 4B, Welch’s *t*-test: *t* = 1.175, *P* = 0.305, df = 4.001).

**Figure 3.**
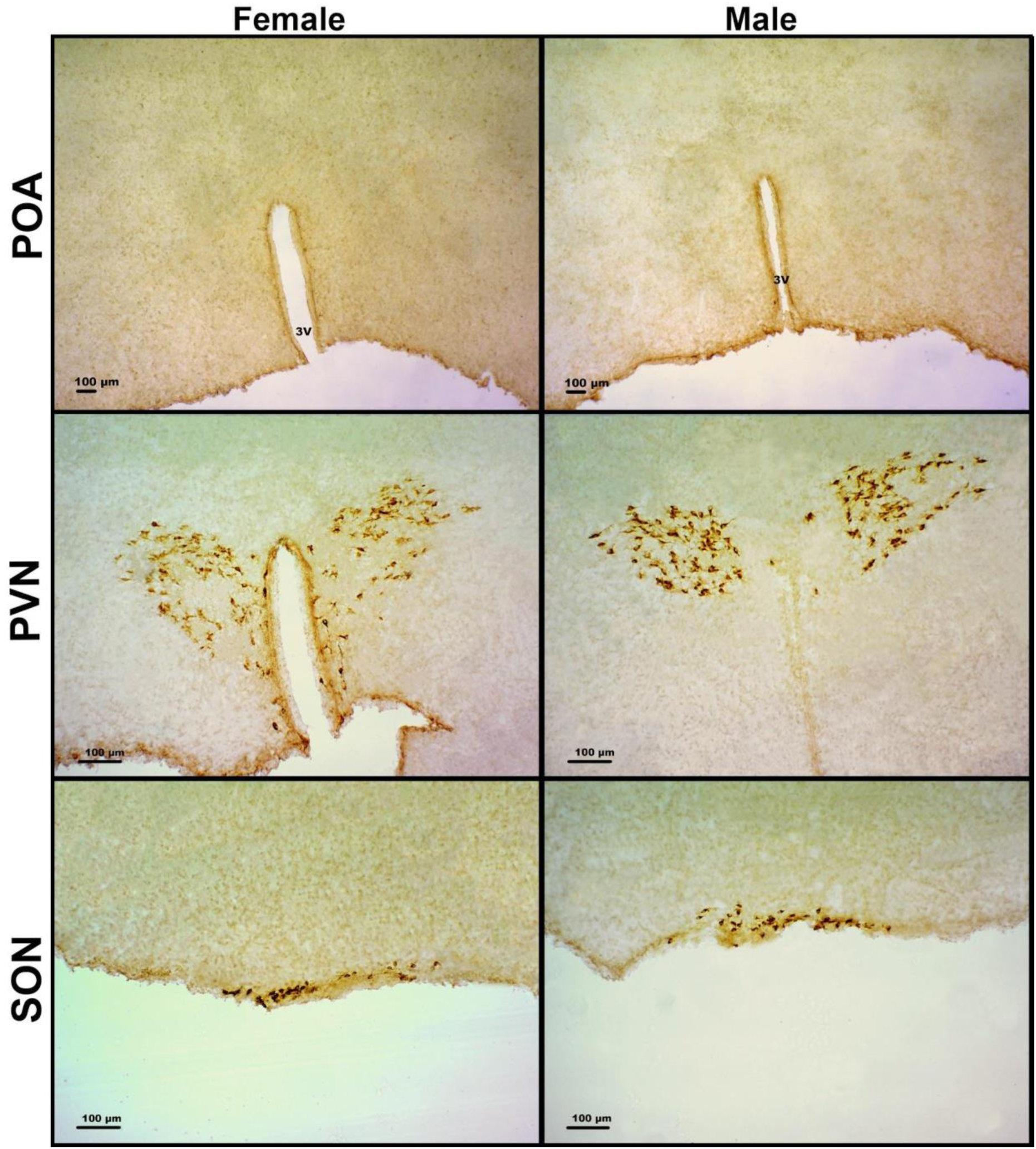
Representative photomicrographs of OXT immunoreactivity in the POA, PVN, and SON of male and female mice at PN7. Representative images of OXT immunostaining in the preoptic area (POA), paraventricular nucleus (PVN), and supraoptic nucleus (SON) were obtained using a Zeiss microscope equipped with a Leica DC200 digital camera. Images of the POA were acquired using a 5x objective, whereas the PVN and SON were acquired using a 10x objective. In each panel, the left image corresponds to a representative female and the right image to a representative male. OXT-immunoreactive neurons and fibers were detected in the PVN, and SON of both sexes. In the POA, OXT-immunoreactive neurons were absent in both males and females.

**Figure 4.**
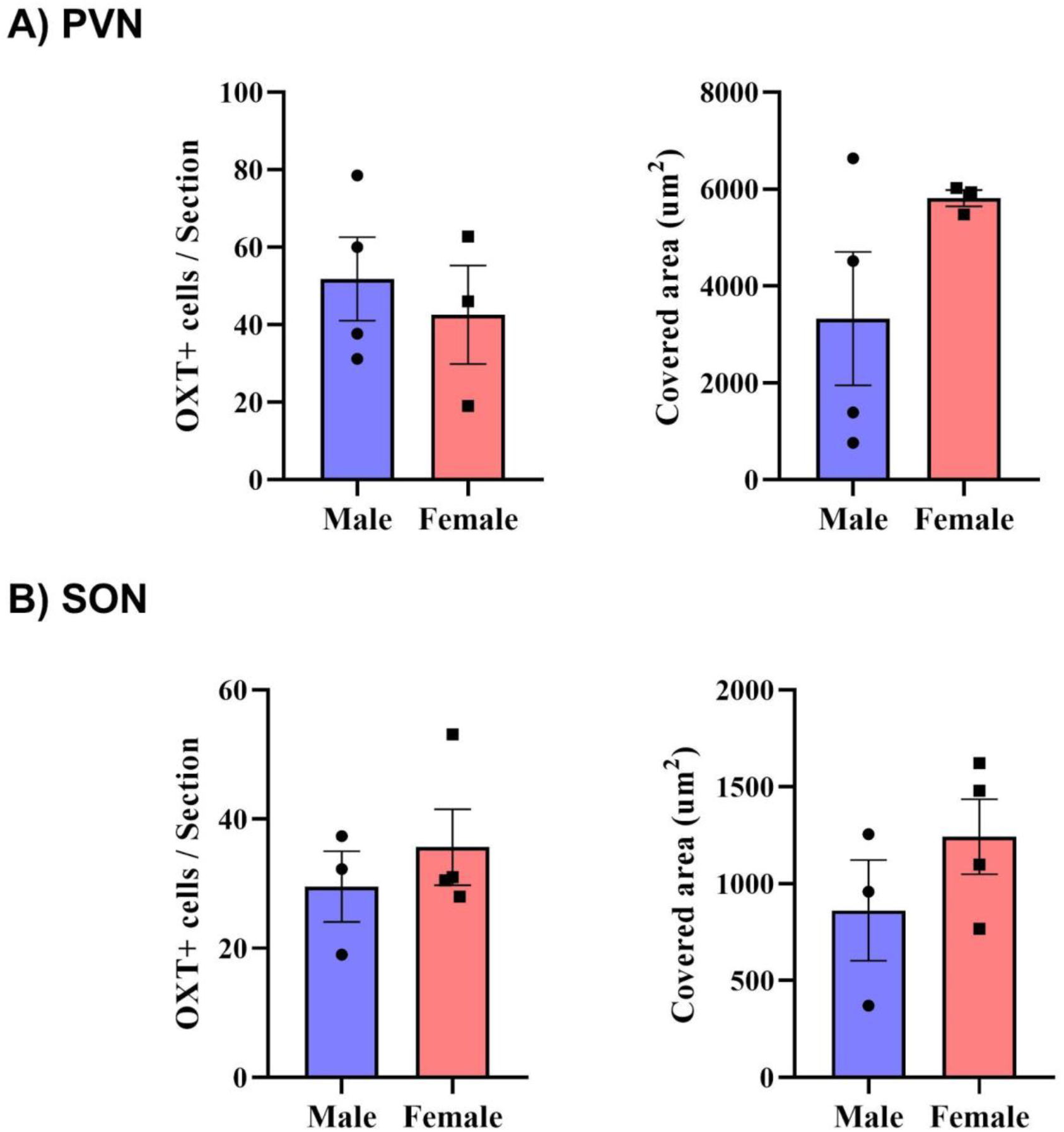
Oxytocin immunoreactivity in the paraventricular and supraoptic nuclei at PN7. Quantification of the total number of OXT-immunoreactive (OXT+) neurons and the percentage of OXT-immunoreactive area in the paraventricular nucleus (PVN; **A**) and supraoptic nucleus (SON; **B**) of male and female mice at postnatal day 7 (PN7). No significant sex differences were detected in either the number of OXT+ neurons or the OXT-immunoreactive area in either brain region. OXT immunoreactivity was not detected in the preoptic area (POA) of either sex. Data are presented as mean ± SEM. Statistical significance was determined using Welch’s *t*-test. *P* < 0.05 was considered statistically significant.

#### 3.3.2 Oxtr mRNA expression at PN7

We next examined whether *Oxtr* mRNA expression exhibited sex-dependent differences during the critical period of brain sexual differentiation. Relative *Oxtr* mRNA expression was quantified in the PFC, POA, and PVN at PN7 by RT-qPCR (Fig. 5). A significant sex difference was detected only in the PFC, where males displayed higher *Oxtr* mRNA levels than females (Fig. 5, Welch’s *t*-test: *t* = 4.302, *P* = 0.006, df = 5.649). In contrast, no significant sex differences were observed in either the POA or the PVN (Fig. 5, Welch’s *t*-test: POA, *t* = 1.409, *P* = 0.209, df = 5.959; PVN, *t* = 0.177, *P* = 0.871, df = 3.036).

**Figure 5.**
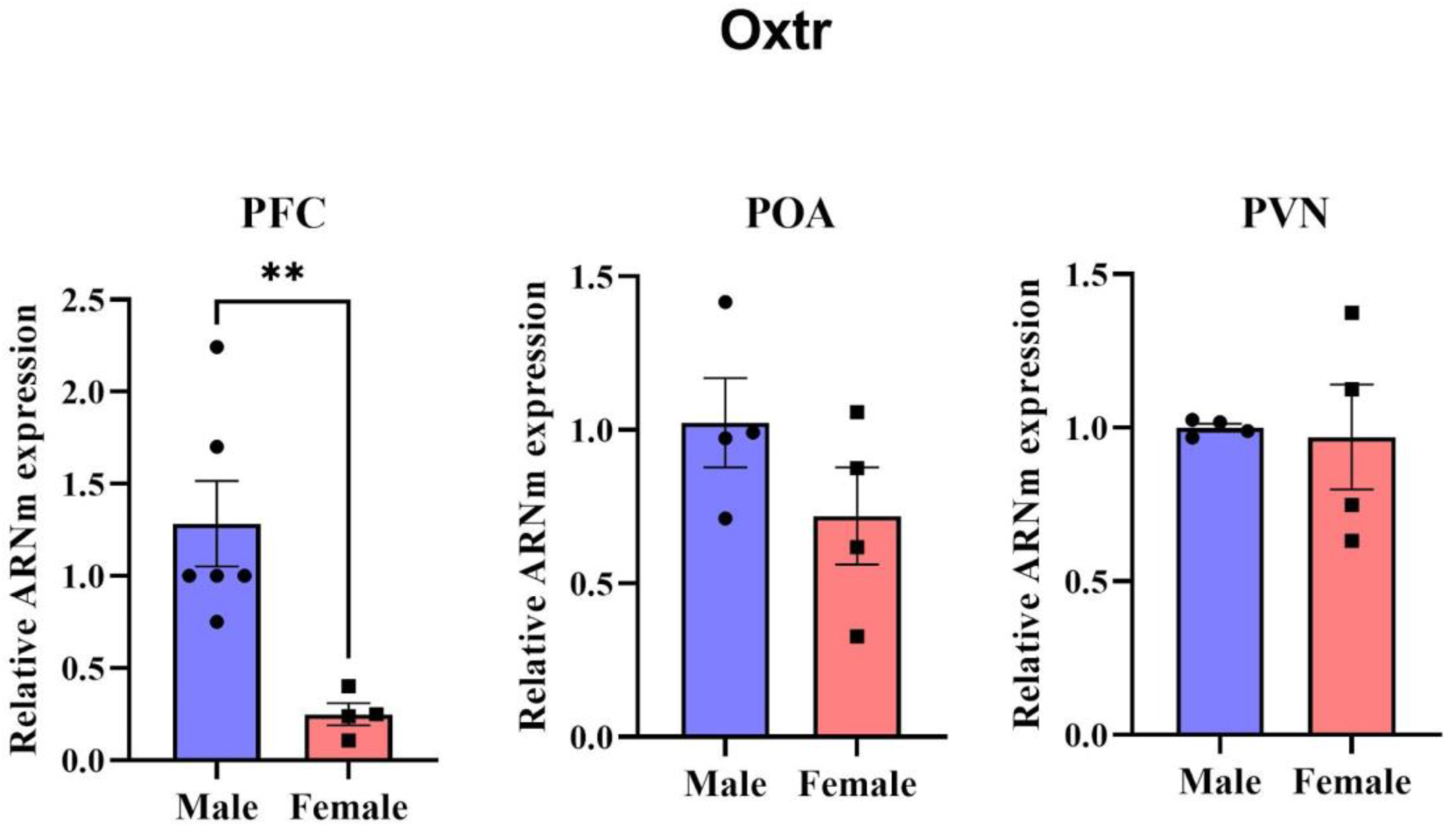
Sex differences in *Oxtr* mRNA expression at PN7. Relative *Oxtr* mRNA expression in the PFC, POA, and PVN of male and female mice at PN7. A significant sex difference was detected only in the PFC, where male mice exhibited higher *Oxtr* mRNA expression than females. No significant sex differences were observed in the POA or PVN. Data are presented as mean ± SEM. Statistical significance was determined using Welch’s *t*-test. *P* < 0.05 was considered statistically significant.

#### 3.3.3 Oxytocin immunoreactivity at PN18

At PN18, OXT-immunoreactive neurons were detected in both the periventricular nucleus (Pe) and the anteroventral periventricular nucleus (AVPe) (Figure 6). Females exhibited a significantly greater number of OXT-immunoreactive neurons than males in the Pe (Fig. 7A, Welch’s *t*-test: *t* = 6.673, *P* = 0.006, df = 3.189), whereas no significant sex differences were observed in the AVPe (Fig. 7B, Welch’s *t*-test: *t* = 0.318, *P* = 0.764, df = 5.000).

**Figure 6.**
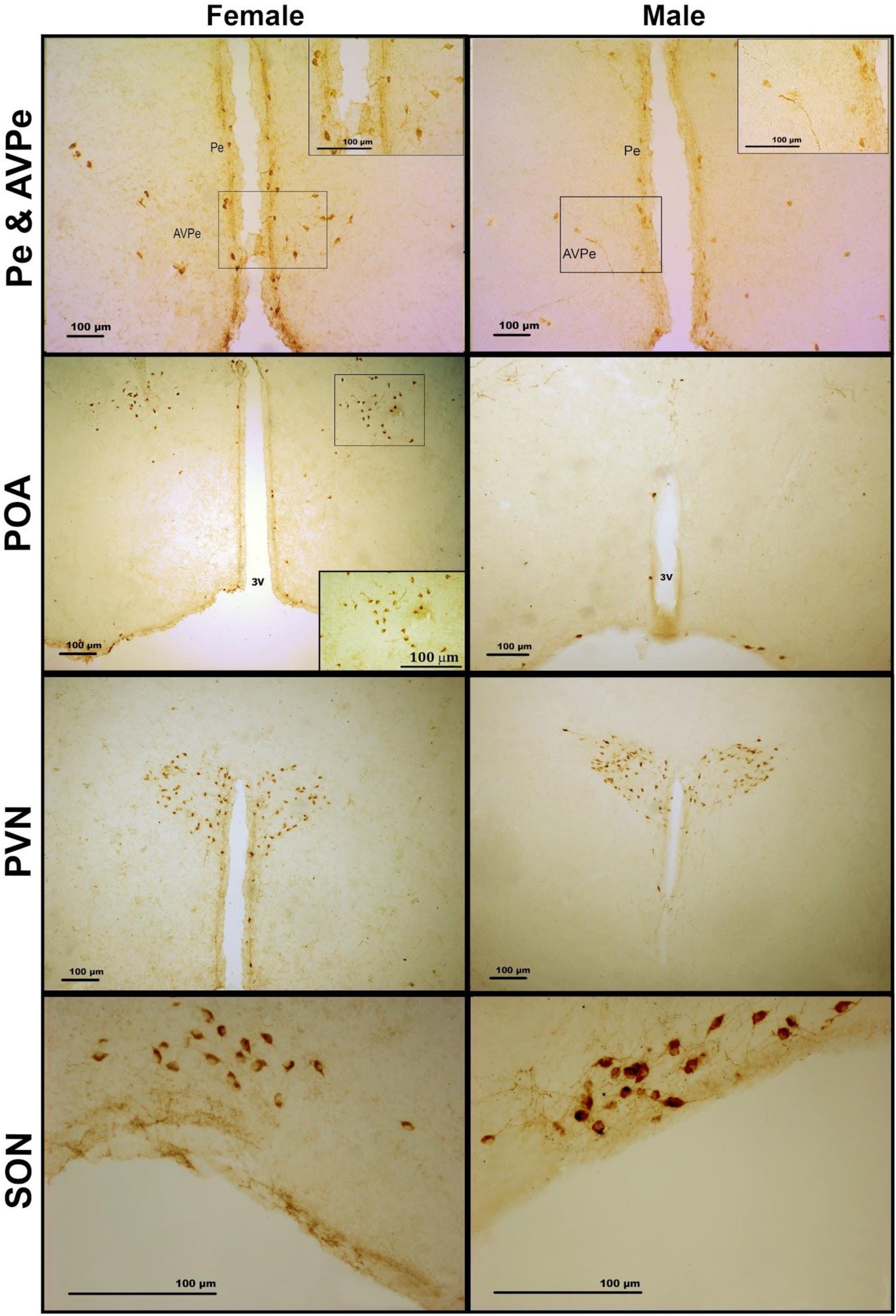
Representative photomicrographs of OXT immunoreactivity in the Pe, AVPe, POA, PVN, and SON of male and female mice at PN18. Representative images of OXT immunostaining in the periventricular nucleus (Pe), anteroventral periventricular nucleus (AVPe), preoptic area (POA, paraventricular nucleus (PVN), and supraoptic nucleus (SON) obtained using a Zeiss microscope equipped with a Leica DC200 digital camera. Images of the Pe, AVPe, POA, and PVN were acquired using a 10x objective, whereas SON images were acquired using a 40x objective. In each panel, the left image corresponds to a representative female and the right image to a representative male. OXT-immunoreactive neurons and fibers were detected in the Pe, AVPe, PVN, and SON of both sexes. In the POA, OXT-immunoreactive neurons were observed in females but were absent in males.

**Figure 7.**
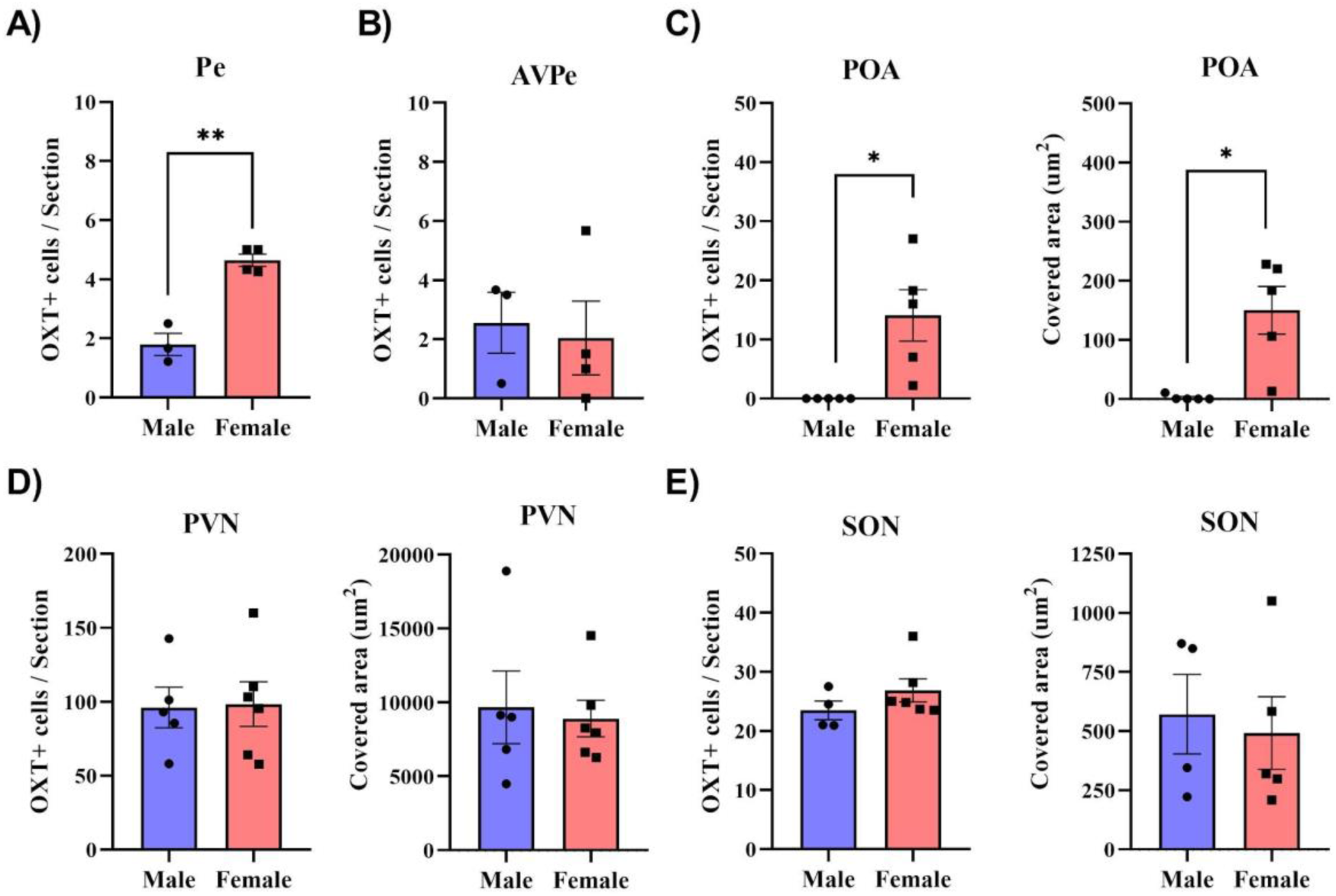
Quantification of OXT immunoreactivity in the Pe, AVPe, POA, PVN, and SON at PN18. Quantification of the total number of OXT-immunoreactive (OXT+) neurons in the periventricular nucleus (Pe; **A**), anteroventral periventricular nucleus (AVPe; **B**), preoptic area (POA; **C**), paraventricular nucleus (PVN; **D**), and supraoptic nucleus (SON; **E**) of male and female mice at PN18. The percentage of OXT-immunoreactive area, reflecting the density of OXT-positive fibers within each region of interest, was also quantified in the POA (C), PVN (D), and SON (E). Female mice exhibited a significantly greater number of OXT+ neurons in the Pe and POA, as well as a greater OXT-immunoreactive area in the POA, compared with males. No significant sex differences were detected in the AVPe, PVN, or SON. Data are presented as mean ± SEM. Statistical significance was determined using Welch’s *t*-test. *P* < 0.05 was considered statistically significant.

In the POA, females also exhibited a significantly greater number of OXT-immunoreactive neurons than males (Fig. 7C, Welch’s *t*-test: *t* = 3.239, *P* = 0.032, df = 4.000). Notably, no OXT-immunoreactive cells were detected in the POA of male mice, whereas females displayed an average of 14.09 OXT-positive neurons. Analysis of the OXT-immunoreactive area, which reflects the density of OXT-positive fibers within the region of interest, also revealed significantly greater values in females than in males (Fig. 7C, Welch’s *t*-test: *t* = 3.646, *P* = 0.022, df = 4.021).

In the PVN and SON, no significant sex differences were observed in either the total number of OXT-immunoreactive neurons or the OXT-immunoreactive area at PN18 (Fig. 7D, E). In the PVN, both the number of OXT-positive neurons (Fig. 7D, Welch’s *t*-test: *t* = 0.114, *P* = 0.912, df = 8.996) and the OXT-immunoreactive area (Fig. 7D, Welch’s *t*-test: *t* = 0.276, *P* = 0.792, df = 5.971) were comparable between males and females. Similarly, no significant sex differences were detected in the SON for either the number of OXT-positive neurons (Fig. 7E, Welch’s *t*-test: *t* = 1.348, *P* = 0.215, df = 7.987) or the OXT-immunoreactive area (Fig. 7E, Welch’s *t*-test: *t* = 0.349, *P* = 0.738, df = 6.616).

#### 3.3.4 Oxtr mRNA expression at PN18

To determine whether the sex difference in *Oxtr* expression observed during the critical period of sexual differentiation persisted after the end of this developmental window, we quantified the relative *Oxtr* mRNA expression in the PFC, POA, and PVN at PN18 (Fig. 8). No significant sex differences in *Oxtr* mRNA expression were detected in any of the brain regions analyzed (Fig. 8, Welch’s t-test: PFC, *t* = 0.170, *P* = 0.872, df = 4.871; POA, *t* = 0.672, *P* = 0.560, df = 2.416; PVN, *t* = 0.539, *P* = 0.611, df = 5.650). These findings indicate that the sex difference in *Oxtr* expression observed in the PFC during the critical period was no longer present by PN18.

**Figure 8.**
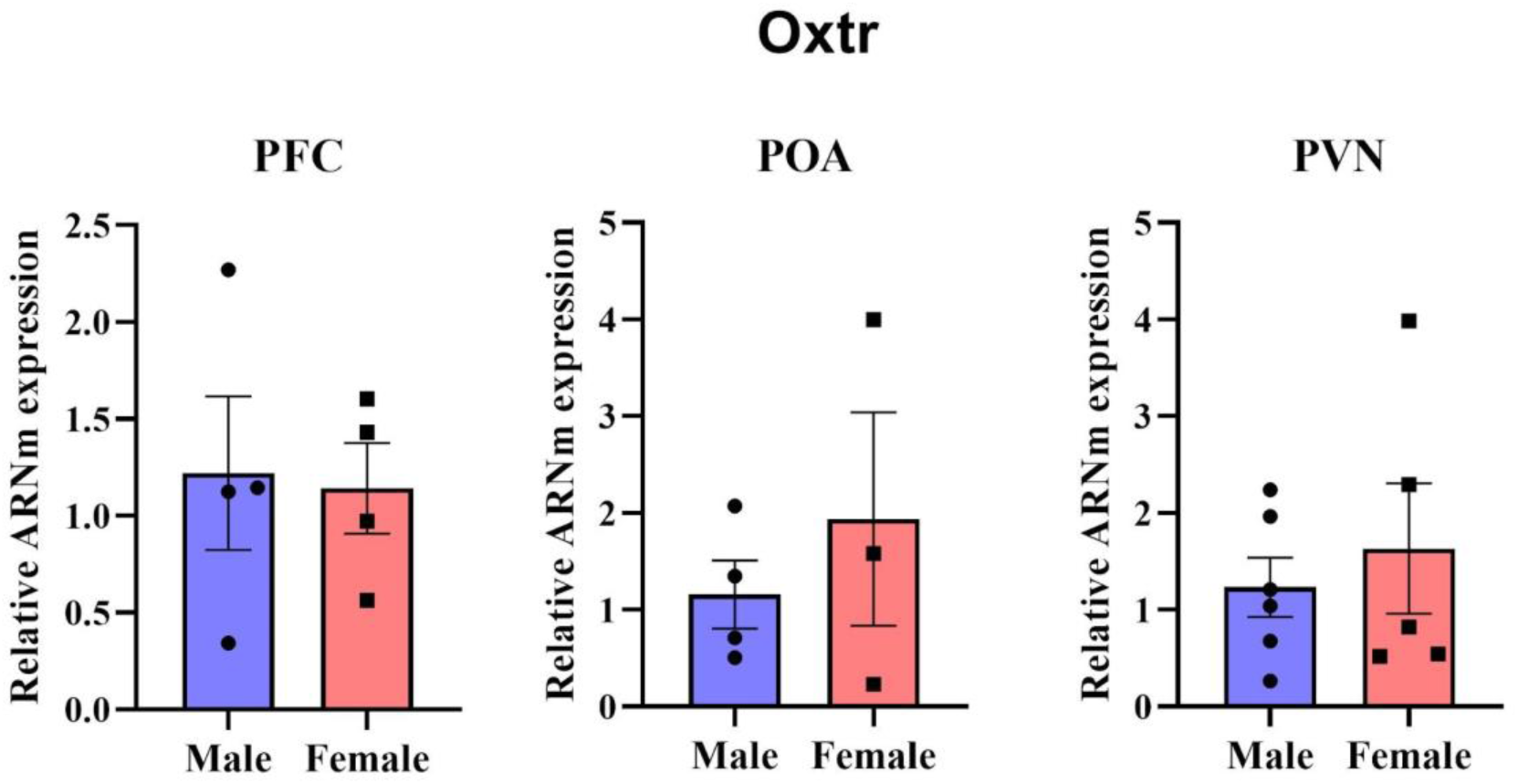
Relative *Oxtr* mRNA expression in the prefrontal cortex, preoptic area, and paraventricular nucleus at PN18. Relative *Oxtr* mRNA expression was quantified by RT-qPCR in the PFC, POA and PVN of male and female mice at PN18. No significant sex differences in *Oxtr* mRNA expression were detected in any of the brain regions analyzed. Data are presented as mean ± SEM. Statistical significance was assessed using Welch’s *t*-test. *P* < 0.05 was considered statistically significant.

## 4. Discussion

In the present study, we found that the expression of genes encoding the active DNA demethylation and repair machinery (*Tet1*, *Tet2*, *Tet3*, *Gadd45a*, *Gadd45b*, *Tdg*) differed between males and females at PN7 specifically in the PFC, while no sex differences in this machinery were detected in the other two regions examined by qPCR (POA and PVN) at either PN7 or PN18. A parallel, region-and age-specific pattern was observed for *Oxtr*: expression was higher in males than in females in the PFC at PN7, with no sex difference detected in the POA or PVN at either age, and no difference in any region, including the PFC, by PN18. In parallel, oxytocin expression was undetectable in the POA, Pe, and AVPe at PN7, but by PN18 it was present in all regions examined (POA, Pe, AVPe, PVN and SON), consistent with previously described developmental profiles of the oxytocinergic system. Notably, we found a sex difference in oxytocin expression, higher in females than males, specifically in the POA and Pe at PN18, a finding that, to our knowledge, has not been previously reported.

The first evidence that active epigenetic mechanisms, rather than passive hormonal absence, orchestrate the sexually differentiated state of the brain was provided by Nugent et al. (2015), who showed that DNA methyltransferase (DNMT) activity is higher in the female POA and actively represses masculinizing genes; a perinatal reduction in DNMT activity, driven by gonadal steroids, releases this repression and permits masculinization. Pharmacological DNMT inhibition or conditional *Dnmt3a* deletion in females was sufficient to masculinize neuronal phenotype and behavior, demonstrating that feminization of the brain must be actively maintained through DNA methylation rather than passively defaulted to. If methylation actively represses the male-typical program, then the enzymatic machinery responsible for actively removing methylation marks would be expected to be preferentially engaged in males, the sex undergoing masculinization during the perinatal sensitive period. Consistent with this, in (Cisternas, Cortes, Bruggeman, et al., 2020) we reported that expression of the *Tet* demethylases in hypothalamic regions of the mouse brain peaks during the first postnatal week and is higher in males than females, with this difference narrowing as development proceeds. The sex difference we observed here in the expression of the active DNA demethylation and repair machinery (*Tet1*, *Tet2*, *Tet3*, *Gadd45a*, *Gadd45b*, *Tdg*) was evident at PN7 specifically in the PFC, and absent, at either age, in any of the other regions examined (POA or PVN). Together, these findings support the view that, where present, active DNA demethylation is preferentially organized during the perinatal critical period of brain sexual differentiation, rather than constituting a stable, lifelong feature of the epigenetic landscape, and that this window of sexually dimorphic epigenetic regulation is itself regionally restricted.

The oxytocinergic system has been previously reported to be regulated by gonadal steroids (Sharma et al., 2019). However, the sex difference we observed in oxytocin expression in the POA and Pe at PN18, higher in females than males, occurs at a developmental stage that precedes the onset of puberty and the resumption of gonadal steroidogenesis in the mouse, making it unlikely to depend on the activational effects of pubertal or reproductive-stage ovarian hormones. Instead, this difference more plausibly reflects the organizational action of the perinatal testosterone surge, which peaks in males around embryonic day 17/18 and again shortly after birth (PN0), and is known to permanently redirect the sexual differentiation of hypothalamic and preoptic circuits (Arnold, 2009). A direct, hormone-independent contribution of the sex chromosome complement (XX vs XY) cannot be ruled out as an additional or complementary mechanism, consistent with the broader unified model of mammalian sexual differentiation, which recognizes gonadal hormone-independent effects of sex chromosome genes alongside the classical organizational/activational hypothesis (Arnold, 2017).

This timing is consistent with independent evidence that the oxytocinergic system undergoes its most dynamic specification precisely during this perinatal-to-early-postnatal window. Using whole-brain 3D reconstructions, Madrigal and Jurado (2021) showed that OXT and vasopressin (AVP) neurons across hypothalamic nuclei display distinctive developmental trajectories and marked cellular plasticity from embryonic to early postnatal stages, with a substantial population of neurons transiently co-expressing OXT and AVP during the first postnatal week before this mixed phenotype declines toward adulthood. Similarly, using a four-timepoint postnatal atlas (P0, P3, P7, P14), Soumier et al. (2022) found that, unlike the AVP system, the OXT system continues to mature well into the postnatal period: OXT neuron number doubles with region-specific dynamics in the periventricular and paraventricular nuclei, and PVN cells gradually acquire an oxytocinergic phenotype over this same window. This dynamic, strain-consistent pattern of oxytocin system maturation is further supported by findings of Hammock and Levitt (2013), who mapped OXTR ligand binding from embryonic tissue through postnatal development in the C57BL/6J mouse, and found that several brain regions display dense, transient OXTR binding during development that is largely absent in the adult brain, underscoring that this receptor system undergoes substantial, region-specific reorganization precisely across the perinatal-to-juvenile window we examined here. Together, these findings indicate that the identity and number of oxytocin-expressing neurons in these nuclei are still being actively established during precisely the period when the sex differences in demethylation/repair machinery reported here are most pronounced, providing a plausible cellular substrate through which perinatal testosterone-driven epigenetic programming could differentially shape OXT expression in males and females.

Convergent evidence for a hypothalamus-specific, early-established sexual dimorphism of the oxytocin system comes from receptor-mapping studies. Newmaster et al. (2020) generated a brain-wide, cellular-resolution atlas of OXTR expression across postnatal development and found that, unlike most brain regions, where OXTR expression is transiently upregulated and then downregulated with age regardless of sex, the hypothalamus retains sexually dimorphic OXTR expression, with denser OXTR labeling in the ventral premammillary nucleus of males emerging as early as P14, well before puberty. In cortical regions, including the prefrontal cortex, both laboratories (Newmaster et al., 2020) and (Hammock & Levitt, 2013) described a transient, non-sex-specific developmental trajectory of OXTR expression, peaking around the second to third postnatal week and declining thereafter. The sex difference we observed in the PFC - higher *Oxtr* expression in males specifically at PN7, resolved by PN18 - extends this picture: it indicates that, transiently and prior to the reported cortical expression peak, *Oxtr* expression in the prefrontal cortex can also be sexually dimorphic, suggesting that sexually dimorphic regulation of the oxytocin system is not strictly confined to the hypothalamus, but can also emerge, earlier and more transiently, in a cortical region relevant to the same critical period of epigenetic regulation described above. Sharma et al. (2019) similarly reported a female-biased number of OXTR-expressing neurons in the medial preoptic area, driven specifically by the anteroventral periventricular nucleus (AVPV); notably, this dimorphism required intact adult ovaries and was abolished by ovariectomy, indicating an activational, estrogen-dependent component in adulthood. The contrast between the activational regulation described by (Sharma et al., 2019) in gonadally intact adults and the pre-pubertal emergence reported by (Newmaster et al., 2020) and in the present study illustrates that *Oxtr*/OXTR dimorphisms can arise through distinct, region-and age-specific mechanisms, some organizational and established early, others requiring ongoing adult hormone signaling. Both sex differences we describe here - the higher *Oxtr* expression in the PFC of males at PN7, and the higher oxytocin expression in the POA and Pe of females at PN18 - occur in neonatal or juvenile neonatal animals, before the pubertal rise in gonadal steroids, and are therefore more consistent with the former: organizational effects of the perinatal testosterone surge, plausibly relayed through the sex-specific epigenetic machinery examined in this study, rather than activational effects of reproductive-stage hormones. Further supporting a region-specific, sex-and age-dependent regulation of the oxytocin system (Vaidyanathan & Hammock, 2020) reported that genetic loss of *Oxtr* in C57BL/6J mice altered Oxytocin mRNA expression specifically in the PVN, but not the SON, following a sex-and age-dependent trajectory already apparent at P14 and persisting, in a sex-specific manner, into adulthood (P90). This regional dissociation between the PVN and the SON parallels the anatomical distinction central to the present study and reinforces the idea that oxytocin expression in the PVN, in particular, is subject to a sex-and developmental stage-specific regulatory program that could plausibly be shaped by the epigenetic mechanisms we describe.

Direct causal support for this model comes from studies manipulating DNA methylation/demethylation during the neonatal period. Pharmacological inhibition of DNA methylation disrupts the testosterone-dependent masculinization of calbindin-expressing neurons in the POA and BNST and reduces the sex difference in ERα-expressing neurons in the VMH, indicating that active methylation turnover is required for gonadal hormones to establish normal sexually dimorphic neurochemical phenotypes (Cisternas, Cortes, Golynker, et al., 2020). Similarly, the sex difference in ERα expression in the arcuate nucleus depends on differential methylation and TET-mediated demethylation of the *Esr1* promoter (Cortes et al., 2022). A causal role specifically for the *Gadd45*-dependent arm of the demethylation pathway examined here is provided by (Stockman et al., 2022), who reported that neonatal male rats generate roughly twice as many new neurons as females in the dentate gyrus, an effect associated with higher global DNA methylation and lower *Gadd45a* expression in the male dentate gyrus at birth; pharmacological inhibition of DNA methylation eliminated both the elevated methylation and the sex difference in cell proliferation in males, with no effect in females, directly linking reduced *Gadd45a*-dependent demethylation to a male-biased developmental outcome within the critical period examined in the present study. Together with (Nugent et al., 2015), these studies demonstrate that the epigenetic machinery examined here - methylation, TET-mediated demethylation, and *Gadd45*-dependent base-excision repair is not merely a passive epigenetic marker of sex, but an active, bidirectional mechanism, methylation restraining and demethylation permitting sexually dimorphic gene expression, lending plausibility to the idea that comparable processes act on *Oxtr* and related genes to shape the sexually dimorphic organization of the oxytocinergic system.

## 5. Conclusion

The present study shows an age-and brain region-specific pattern of expression of the active DNA demethylation machinery in male and female mice. Sex differences in this machinery were detected only during the first postnatal week, and only in the prefrontal cortex, where expression was higher in males; this difference was no longer present by PN18.

A parallel, region-and age-specific pattern was found for *Oxtr*, with higher expression in males than in females in the prefrontal cortex at PN7, and no sex difference in the other two regions examined by qPCR (POA and PVN) at either age. Oxytocin itself was undetectable in the POA, Pe, and AVPe at PN7, but by PN18 was present in all regions examined (POA, Pe, AVPe, PVN and SON), with a clear sex difference in the POA and Pe, higher in females than males. Together, these findings reveal a region-and age-specific pattern of sexual dimorphism, in which sex differences in the DNA demethylation/repair machinery and in *Oxtr* are jointly confined to the prefrontal cortex during the earliest postnatal window examined, whereas sex differences in oxytocin peptide expression emerge later in development and are restricted to discrete hypothalamic/preoptic nuclei once its expression becomes detectable. Together with the mechanistic evidence reviewed above, these results support a model in which the sex-specific distribution of active DNA demethylation and repair machinery during discrete postnatal windows may contribute to the organizational, testosterone-dependent shaping of the oxytocinergic system relevant to social behavior.

## AUTHOR CONTRIBUTIONS

**C. D. Cisternas** conceived and designed the research. **R. Bigarani**, **B. Ghione** and **C.D. Cisternas** performed all experiments and analyzed data. RB, BG, MJC and CDC interpreted the results of the experiments. **R. Bigarani** and **B. Ghione** elaborated figures. **R. Bigarani** and **C. D. Cisternas** wrote the first draft of the manuscript. **C.D. Cisternas** and **M. J. Cambiasso** were responsible for resources, project administration, and funding acquisition, as well as editing and revising the manuscript with critical input from all authors. All authors read and approved the final manuscript. **C. D. Cisternas** drafted the manuscript.

## ACKNOWLEDGEMENTS

We gratefully acknowledge the staff of the INIMEC animal facility and Dr. Soledad de Olmos for their dedicated technical support. The authors are grateful to Dr. Andrea Godino for feedback on the manuscript. The authors acknowledge the use of Claude and ChatGPT AI tools to assist grammar and language improvement of text based on author-directed concepts. These tools were not used autonomously, and all outputs were reviewed and validated by the authors. The authors assume full responsibility for the accuracy, integrity, and originality of the work. **C.D. Cisternas** and **M. J. Cambiasso** are researchers of CONICET-Argentina.

## ETHICS APPROVAL STATEMENT

All procedures were approved by the Institutional Committee for the Care and Use of Experimental Animals (CICUAL) of Instituto Ferreyra (Institutional Resolution 1/2020) and conducted in accordance with national and international guidelines for the care and use of laboratory animals.

## DATA AVAILABILITY STATEMENT

Data will be available upon request to the corresponding author, Carla Cisternas

