## Supplementary material for "Sex differences in DNA demethylation machinery precede sex differences in the oxytocinergic system in the postnatal mouse brain": Suplemental material

**Supplemental material**

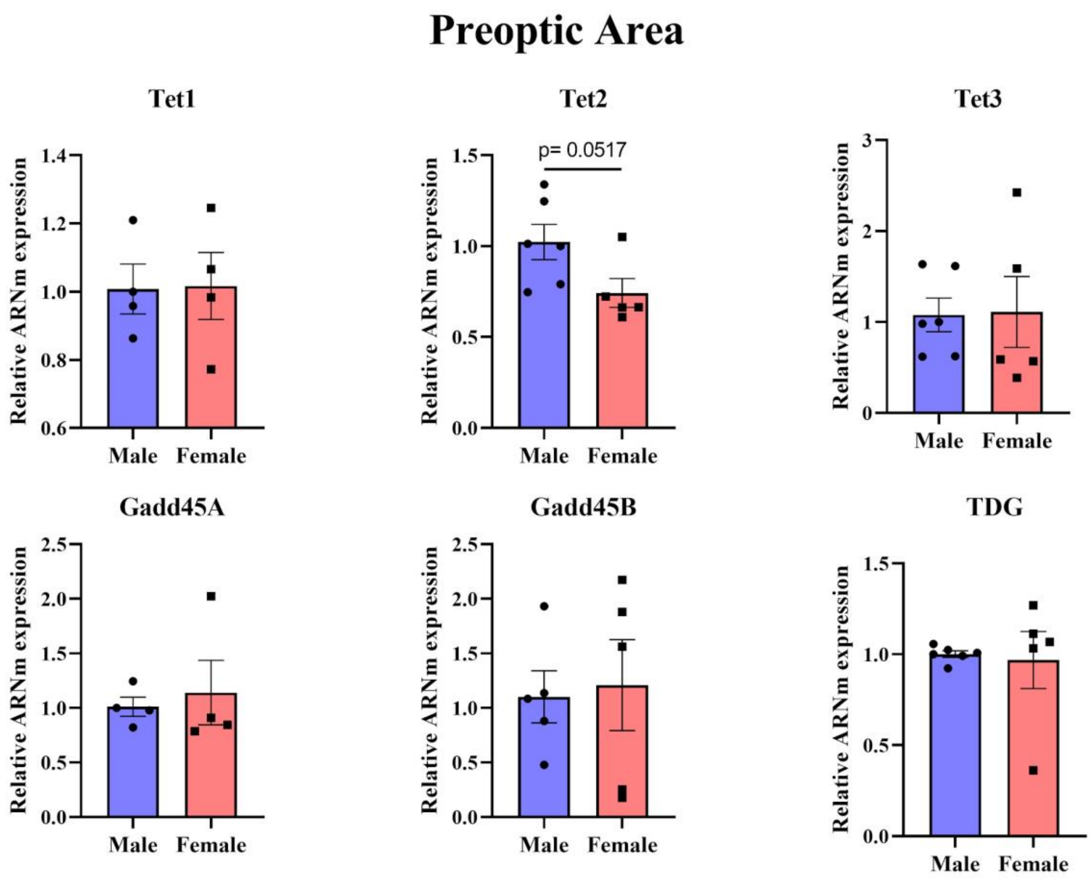


**Supplemental Figure 1. Evaluation of sex differences in the expression of active DNA demethylation and DNA repair genes in the preoptic area at PN7.** Relative mRNA expression of the active DNA demethylation enzymes *Tet1*, *Tet2*, and *Tet3*, and the DNA repair-associated genes *Gadd45a*, *Gadd45b*, and *Tdg* in the POA of male and female mice at PN7. No significant sex differences were detected for any of the genes analyzed, although *Tet2* expression showed a trend toward higher mRNA levels in males compared with females. Data are presented as mean ± SEM. Statistical significance was determined using Welch's *t*-test. $P < 0.05$ was considered statistically significant.

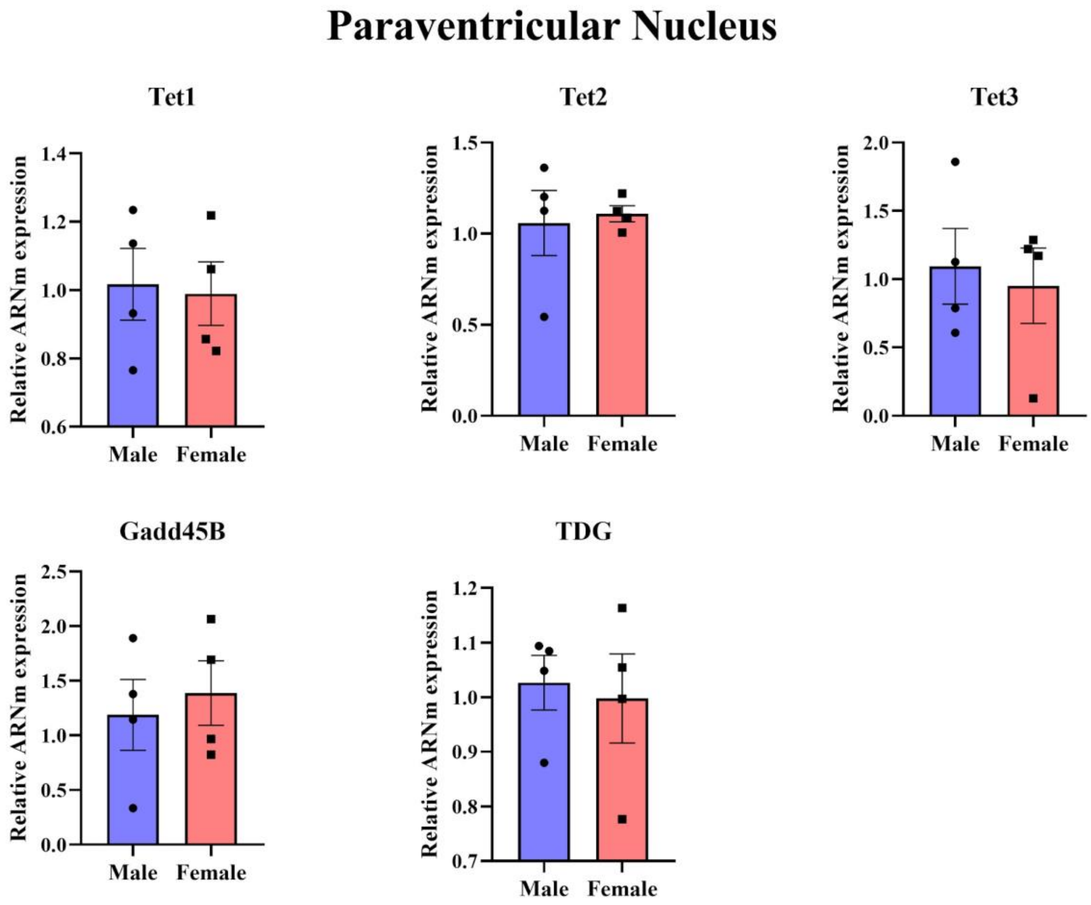


**Supplemental Figure 2. Evaluation of sex differences in the expression of active DNA demethylation and DNA repair genes in the paraventricular nucleus at PN7.** Relative mRNA expression of the active DNA demethylation enzymes *Tet1*, *Tet2,* and *Tet3,* and the DNA repair-associated genes *Gadd45b* and *Tdg* in the PVN of male and female mice at PN7. No significant sex differences were detected for any of the genes analyzed. *Gadd45a* mRNA expression was not detected in this region at PN7. Data are presented as mean ± SEM. Statistical significance was determined using Welch's *t*-test. $P < 0.05$ was considered statistically significant.

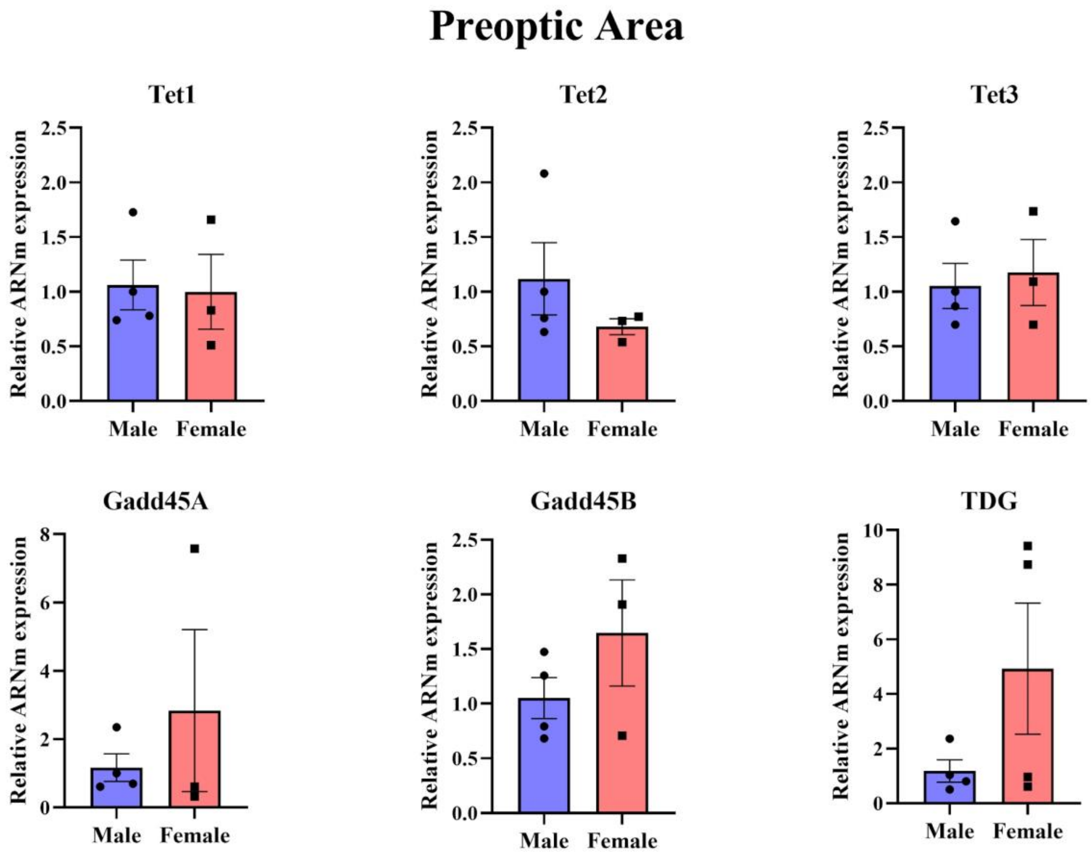


**Supplemental Figure 3. Expression of active DNA demethylation and DNA repair genes in the preoptic area at PN18.** Relative mRNA expression of the active DNA demethylation enzymes *Tet1*, *Tet2*, and *Tet3*, and the DNA repair-associated genes *Gadd45a*, *Gadd45b*, and *Tdg* in the POA of male and female mice at PN18. No significant sex differences were detected for any of the genes analyzed. Data are presented as mean ± SEM. Statistical significance was determined using Welch's *t*-test. $P < 0.05$ was considered statistically significant.

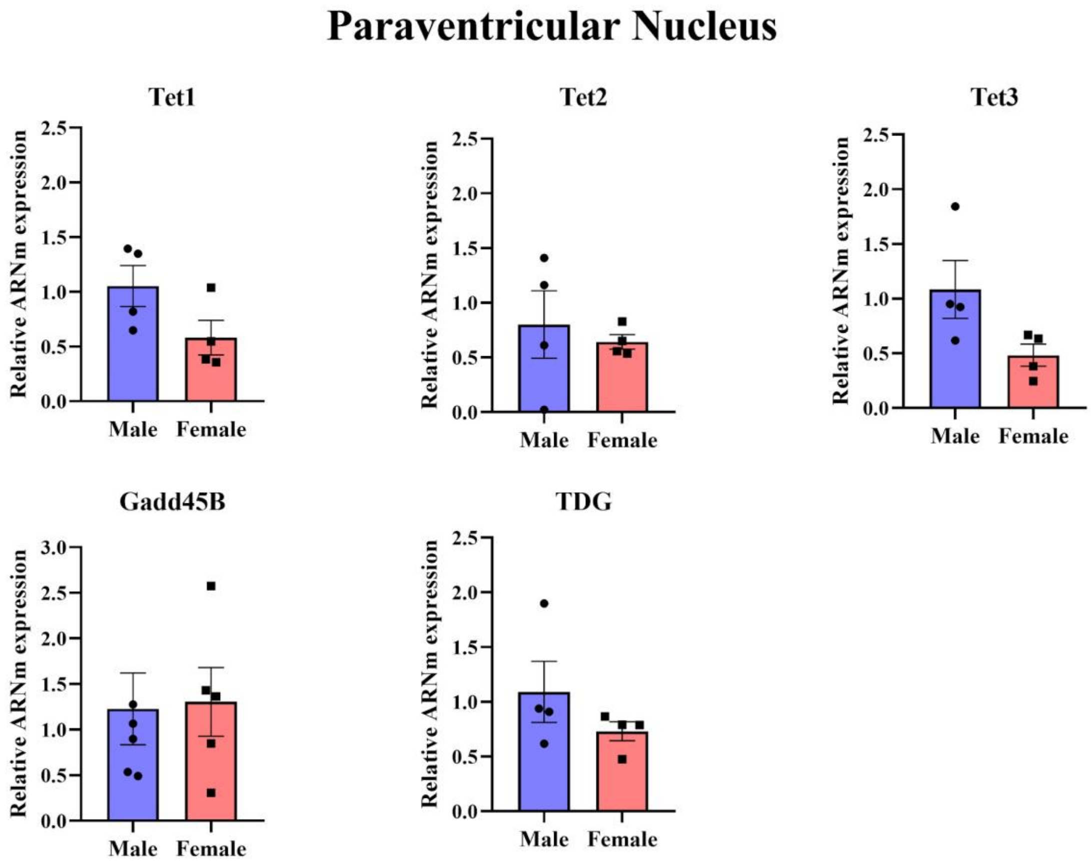


**Supplemental Figure 4. Expression of active DNA demethylation and DNA repair genes in the paraventricular nucleus at PN18.** Relative mRNA expression of the active DNA demethylation enzymes *Tet1*, *Tet2*, and *Tet3*, and the DNA repair-associated genes *Gadd45b* and *Tdg* in the PVN of male and female mice at PN18. No significant sex differences were detected for any of the genes analyzed. *Gadd45a* mRNA expression was not detected in this region. Data are presented as mean ± SEM. Statistical significance was determined using Welch's *t*-test. $P < 0.05$ was considered statistically significant.
